# Moonlighting on the *Fasciola hepatica* tegument: enolase, a glycolytic enzyme, interacts with the extracellular matrix and fibrinolytic system of the host

**DOI:** 10.1101/2024.03.15.585155

**Authors:** Eve O’Kelly, Krystyna Cwiklinski, Carolina De Marco Verissimo, Nichola Eliza Davies Calvani, Jesús López Corrales, Heather Jewhurst, Andrew Flaus, Richard Lalor, Judit Serrat, John P. Dalton, Javier González-Miguel

## Abstract

Enolase is a 47 kDa enzyme that functions within the glycolysis and gluconeogenesis pathways involved in the reversible conversion of D-2-phosphoglycerate (2PGA) to phosphoenolpyruvate (PEP). However, in the context of host-pathogen interactions, enolase from different species of parasites, fungi and bacteria have been shown to contribute to adhesion processes by binding to proteins of the host extracellular matrix (ECM), such as fibronectin (FN) or laminin (LM). In addition, enolase is a plasminogen (PLG)-binding protein and induces its activation to plasmin, the main protease of the host fibrinolytic system. These secondary ‘moonlighting’ functions of enolase are suggested to facilitate pathogen migration through host tissues. This study aims to uncover the moonlighting role of enolase from the parasite *Fasciola hepatica*, shedding light on its relevance to host-parasite interactions in fasciolosis, a global zoonotic disease of increasing concern. A purified recombinant form of *F. hepatica* enolase (rFhENO), functioning as an active homodimeric glycolytic enzyme of ∼94 kDa, was successfully obtained, fulfilling its canonical role. Immunoblotting studies on adult worm extracts showed that the enzyme is present in the tegument and the excretory/secretory products of the parasite, which supports its key role at the host-parasite interface. Confocal immunolocalisation studies of the protein in newly excysted juveniles and adult worms also localised its expression within the parasite tegument. Finally, we showed by ELISA that rFhENO can act as a parasitic adhesin by binding host LM, but not FN. rFhENO also binds PLG and enhances its conversion to plasmin in the presence of the tissue-type and urokinase-type PLG activators (t-PA and u-PA). This moonlighting adhesion-like function of the glycolytic protein enolase could contribute to the mechanisms by which *F. hepatica* efficiently invades and migrates within its host and encourages further research efforts that are designed to impediment this function by vaccination or drug design.

**AUTHOR SUMMARY:** *Fasciola hepatica* is a parasitic worm causing fasciolosis, primarily affecting herbivorous mammals and posing a significant veterinary problem. Furthermore, it is a zoonosis, meaning it can be transmitted to humans. *F. hepatica* enters the definitive host through ingestion of contaminated aquatic plants, migrating through the intestine to settle in the liver bile ducts, where it matures into the adult stage. To migrate, it utilizes various invasion strategies, including the use of multifunctional proteins, known as ‘moonlighting’. In this study, we produced and molecularly characterized the parasitic enzyme enolase as a moonlighting protein to understand *F. hepatica* invasion mechanisms. We produced recombinant enolase with glycolytic activity, its canonical function in parasite energy production. Additionally, we localised this enzyme in the parasite’s tegument, in direct contact with the host, and demonstrated its ability to elicit an immune response early in ovine infection. Finally, we demonstrated the ability of enolase to interact with the extracellular matrix and the host’s fibrinolysis, a proteolytic system responsible for dissolving blood clots. These secondary functions of *F. hepatica* enolase, described here for the first time, along with its localisation and immunogenicity, suggest this protein as an interesting antigen for fasciolosis diagnosis and/or control.

## 1. INTRODUCTION

Despite the obvious energy cost implications, there are a considerable number of helminth parasites whose lifecycles require migration routes from one tissue of the host to another. In fact, it has been suggested that migration is a selectively advantageous strategy in terms of development, host immune-defence and, hence, parasite survival [1,2]. It has been postulated that to successfully invade host tissues, pathogens must first establish contact with the host extracellular matrix (ECM) by highly specific adhesins [3]. In this sense, pathogen adhesion to ECM proteins, such as laminin (LM) and fibronectin (FN), major components of the basement membrane and interstitial matrix, respectively, has been widely described as one of the initiation mechanisms of the infection process [4,5]. Once adhered, the invasion of these physical barriers relies on the secreted proteolytic machinery of helminth parasites, often involving an array of proteases with different peptide bond specificity [6–8]. Additionally, helminth parasites co-opt the proteolytic functions of the host, such as the one exerted by the fibrinolytic system, to invade the host tissues more efficiently. One such example is the expression of proteins with lysine-rich motifs on their surfaces that bind to host plasminogen (PLG), the main fibrinolytic zymogen [9]. Bound PLG can then be converted into its active form, plasmin, by the tissue-type or urokinase-type PLG activators (t-PA and u-PA, respectively) [10]. Plasmin is a broad-spectrum protease whose substrates not only include fibrin that forms blood clots, but also proteins from ECM, the complement system, and immunoglobulins [11]. Therefore, the recruitment of the host PLG/plasmin protease system to the surface of migrating parasites could be of paramount importance for the parasite in terms of immune evasion and migration by degrading different components of the host defence systems and physical barriers.

It is not uncommon that ECM adhesion and PLG-binding functions are carried out by proteins on the host-parasite interface that have other canonical functions in intracellular localisations. A striking example of these ‘moonlighting’ parasite proteins is represented by the glycolytic enzyme enolase [12], whose multifunctionality has been highlighted by its role in pathophysiological conditions [13,14] and as a potential parasite virulence factor [15]. Additionally, enolase has been reported as a PLG-binding receptor of helminth parasites, and its role as a potential pathogen adhesin has also been noted [16–20].

Migration mechanisms are of particular significance in the life cycle of the liver fluke *Fasciola hepatica*, a parasitic trematode responsible for fasciolosis in animals and humans worldwide. Following ingestion of metacercariae by the mammalian host, and release of newly excysted juveniles (FhNEJ) in the duodenum, *F. hepatica* undergoes a complex migratory route through the intestinal wall and in the peritoneum before invading the hepatic tissue. Finally, after some weeks burrowing through and feeding on the hepatic parenchyma, *F. hepatica* juveniles gain access to the major biliary ducts, where they mature into adult worms [21,22]. Fasciolosis is considered a major zoonotic threat, with up to 180 million people at risk of infection, and a serious problem for livestock worldwide, causing high economic losses (∼US$3000 million per year) [23,24]. A deeper understanding of host-parasite molecular interactions in fasciolosis is still needed to underpin the discovery and design of new pharmacological or vaccine interventions [25].

Recently, we have described how *F. hepatica* interacts with host ECM and the fibrinolytic system at the intestinal level as a potential strategy used by the parasite to initiate the migration process [26,27]. Nevertheless, further research is needed to define and characterize the parasitic molecules involved in these adhesin-like mechanisms. In the present work, we performed a functional characterization of *F. hepatica* enolase that clarifies its role as a moonlighting protein capable of interacting with the ECM and fibrinolytic system of the host and activating PLG to plasmin on the surface of the parasite.

## 2. MATERIALS AND METHODS

### 2.1. Confirmation of the *F. hepatica* enolase nucleotide sequence

The DNA sequence of *F. hepatica* enolase was identified by BLAST analysis using the sequence previously reported by Davis et al. [28] (uniprot identifier: Q27655) against the *F. hepatica* genome (WormBase ParaSite: PRJEB6687 and PRJEB25283; maker-scaffold10x_935_pilon-snap-gene-0.41) [29]. The sequence was confirmed by PCR amplification. Total RNA was extracted from *F. hepatica* adult parasites using the miRNeasy Mini Kit (Qiagen) as previously described [30] and cDNA synthesized using the High-Capacity cDNA Reverse Transcription Kit (Thermo Fisher Scientific), according to the manufacturer’s instructions. PCR reactions were performed in a reaction volume of 50 μL, containing 2 μL cDNA, 2× DreamTaq PCR Master Mix (Thermo Fisher Scientific) and 1 nM final concentration of forward and reverse primers. Primers were designed to the regions encoding the putative start and stop codons, respectively, based on the in-silico analysis (Q27655: forward primer 5’-ATGGCAATAAAAGCGATCCACGC-3’, reverse primer 5’-TTATGGTCGGCGGAAGTTCTCC-3’; maker-scaffold10x_935_pilon-snap-gene-0.41: forward primer 5’-ATGGCAACGCTCCGCGCCAG-3’, reverse primer 5’-TTATGGTCGGCGGAAGTTCTCC-3’). The PCR cycling conditions consisted of initial denaturation at 95°C for 3 min, followed by 30 cycles at 95°C for 30 sec, 52°C for 30 sec, 72°C for 1 min, and a final extension at 72°C for 10 min. PCR products were analysed by gel electrophoresis and positive products were purified using the NucleoSpin Gel and PCR clean up kit (Clontech, Takara Bio) before being sent for sequencing by Eurofins Genomics (Germany). The resulting sequences were compared with the in-silico predicted gene sequences by alignment using Clustal Omega [31].

### 2.2. Recombinant expression and purification of *F. hepatica* enolase

The *F. hepatica* enolase (FhENO) sequence with the addition of a 6x His-tag at the C-terminus was synthesised and codon optimized for recombinant expression in *Escherichia coli*. The optimized sequence was cloned into a pET-28a (+) vector containing a kanamycin resistant gene (Genscript) and transformed into *E. coli* BL21 ClearColi cells (DE3; Thermo Fisher Scientific) by electroporation. Transformed cells were selected after growth on Luria Broth Miller (LBM)-Agar supplemented with 50 µg/mL kanamycin and grown overnight at 37°C.

The transformed cells were grown overnight in 50 mL LBM-broth supplemented with kanamycin at 37°C rotating at 160 rpm and then inoculated into LBM-broth with 50 µg/mL kanamycin and grown until an OD_600_ of 0.5 was reached. Expression of rFhENO was induced via the addition of isopropyl β-D-1-thiogalactopyranoside (IPTG) at a concentration of 0.2 mM followed by incubation at 30°C for four hours. The resultant culture was centrifuged at a speed of 10,000 × *g* for 10 min, the pellet resuspended in ST buffer (10 mM Tris, 150 mM NaCl, pH 8.0) and frozen at -20°C. The cell pellet was thawed on ice, digested with lysozyme at a concentration of 10 µg/mL (Sigma-Aldrich) for 30 min and, after the addition of 0.5% sarkosyl (*w/v*) (Sigma-Aldrich), was sonicated for 6 × 10 seconds at 70% amplitude on ice. The pellet was then centrifuged at 15,000 × *g* for 30 min at 4°C. The supernatant was collected, diluted 1:4 with lysis buffer (50 mM NaH_2_PO_4_, 300 mM NaCl, 5 mM imidazole, pH 8.0) and run over a 500 uL bed volume of NiNTA-Agarose beads (Qiagen). The beads were washed three times with 10 mL of wash buffer (50 mM NaH_2_PO_4_, 300 mM NaCl, 10 mM imidazole, pH 8.0) and rFhENO was recovered with elution buffer (50 mM NaH_2_PO_4_, 300 mM NaCl, 250 mM imidazole, pH 7.0). The eluted protein was dialyzed against phosphate-buffered saline (PBS) overnight in 3 kDa dialysis tubing at 4°C, aliquoted, and stored at -80°C.

The concentration of purified rFhENO was determined using a Bradford assay (Bio-Rad) and purity was determined using SDS-PAGE on 4-20% Mini-PROTEAN® TGX™ Precast Protein Gels (Bio-Rad) stained with Biosafe Coomassie (Bio-Rad). Further confirmation of purification of the recombinant protein was obtained by immunoblots probed with mouse anti-polyhistidine IgG (Sigma-Aldrich) and goat anti-mouse alkaline phosphatase-conjugated antibodies (Thermo Fisher Scientific). Gels and Western blots were imaged using the G:BOX Chemi XRQ imager (Syngene).

*F. hepatica* glyceraldehyde-3-phosphate dehydrogenase (FhGAPDH2; maker-scaffold10x_2706_pilon-snap-gene-0.16) was produced and purified as a recombinant, using the same protocol as outlined above for rFhENO using the *E. coli* BL21 ClearColi cell expression system and purified by NiNTA-Agarose beads (Qiagen).

### 2.3. Size-exclusion chromatography of rFhENO

The tertiary molecular size of rFhENO was established by size exclusion chromatography (SEC) using the Superdex 75 10/300 GL (Tricorn) column at a flow rate of 0.4 mL/min. Three proteins of known molecular weight (MW) were used as markers, namely carbonic anhydrase (29 kDa), bovine serum albumin (66 kDa), and alcohol dehydrogenase (150 kDa) (Sigma-Aldrich). rFhENO (100 µg) was injected onto the column and fractions (200 uL) collected into a 96-well chimney plate (Fisher Scientific).

### 2.4. Enzymatic activity assay of rFhENO

The enzymatic activity of rFhENO fractions purified by affinity chromatography and separated by SEC was assessed by a fluorogenic assay using the commercial Enolase Activity Assay Kit (MAK178; Sigma-Aldrich). The molar concentration of protein in each SEC fraction was determined using the Beer Lambert Law (A = ɛcl) based on the mAU readout at 280nm, which was given for each fraction produced by SEC. The SEC readout identified two major peaks, namely peak 1 containing fractions B4-C1 and peak 2 containing fractions C7-D1 that were pooled and assayed for enzymatic activity of rFhENO calculated at 0.1 µM. The activity was read at λex = 540nm / λem = 590nm in a 96-well plate using a PolarStar Omega Spectrophotometer (BMG LabTech), for 30 min at 37°C. The rFhENO activity was expressed as nmole of H_2_O_2_ generated / µM of rFhENO.

### 2.5. *F. hepatica* extracts

Adult parasites required for somatic protein extraction and recovery of the *F. hepatica* excretory/secretory (FhES) products were sourced from two experimental *F. hepatica* infection studies. Somatic extracts: adult parasites were recovered from sheep were experimentally infected with 120 metacercariae (Aberystwyth isolate; Ridgeway Research Ltd, UK) for 16 weeks, at Teagasc Athenry (Ireland). Experimental procedures at Teagasc were carried out under license from Health Products Regulatory Authority (HPRA) by the EU Directive 2010/63/EU (License No. AE19132/P115), after ethical review by the Teagasc Animal Ethics Committee (TAEC2021-298). FhES products: Adult flukes were recovered sheep experimentally infected with 150 *F. hepatica* metacercariae (Italian isolate; Ridgeway Research Ltd), as previously described by Cwiklinski et al. [32].

Somatic extracts of adult *F. hepatica* (FhSoma) were prepared by homogenising the adult fluke in PBS supplemented with a protease and phosphatase inhibitor cocktail (Sigma-Aldrich, two flukes per mL) using a tissue grinder apparatus before centrifugation at 14,000 × *g* for 20 min at 4°C to remove any cellular debris. *F. hepatica* tegument (FhTeg) extracts were prepared by incubating adult parasites in PBS, 1% Tween 20 (*v/v*), pH 7.4 (two flukes per mL) for 1 hour at 37°C. Following the incubation, the supernatant was separated and centrifuged at 14,000 × *g* for 20 min at 4°C to pellet cellular debris. FhES products were prepared as previously described [33]. Briefly, adult flukes were washed in sterile saline, transferred into T75 tissue culture flasks (Sarstedt) and cultured (one fluke per two mL) in RPMI medium (Gibco) supplemented with 0.1% glucose, 100 U penicillin and 100 mg/mL streptomycin (Gibco), at 37°C and 5% CO_2_. After four hours, the medium was collected and centrifuged at 700 × *g* for 30 min at 4°C.

Finally, the supernatants containing the three parasite extracts (FhSoma, FhTeg and FhES) were collected separately, their protein concentration determined using a Bradford assay, and the aliquots stored at -80°C until use.

### 2.6. Size-exclusion chromatography of FhES

FhES was thawed overnight at 4°C, concentrated using a 3 kDa cut-off concentrator spin column (Pierce) and centrifuged at 10,000 × *g* for 20 min. Concentrated FhES (10 mg total protein) was fractionated (2 mL/fraction) by SEC on a Sephacryl S-200 HR 26/60 column (GE Healthcare) connected to an AKTA Start automated liquid chromatography system (GE Healthcare). The column was calibrated using an LMW gel filtration calibration Kit (GE Healthcare). The fractions containing native FhENO activity (fFhES) (determined as described above), which eluted at approximately 95 kDa, and were concentrated using 3 kDa cut-off concentrator spin columns (Pierce). Protein concentration was determined by BCA Protein Assay Kit (Pierce) prior to analysis by immunoblotting.

### 2.7. Immunoblotting

FhSoma, FhTeg, FhES, and fFhES were resolved by SDS-PAGE on 4-20% Precast Protein Gels (Bio-Rad) at a concentration of 2 µg/lane. The proteins were electro-transferred into nitrocellulose membranes using the Trans-Blot Turbo Transfer System (Bio-Rad) and blocked overnight at 4°C in blocking buffer (5% milk powder (*w/v*), PBS containing 0.05% Tween 20 (*v/v*), PBST). Immunoblots were washed three times with PBST and then probed with either a 1:1,000 dilution of polyclonal rabbit anti-rFhENO (Davids Biotechnologie GmbH) or a 1:1,000 dilution of control pre-immune rabbit sera (Davids Biotechnologie GmbH), for one hour at RT. After washing three times with PBST, the blots were incubated with the secondary antibody, goat anti-rabbit IgG HRP-conjugated (Sigma-Aldrich) at a 1:15,000 dilution for one hour at RT. After a further three washes, the immunoblots were developed with SigmaFast 3,3’-diamino-benzidine substrate (Sigma-Aldrich) and imaged using the G:BOX Chemi XRQ imager (Syngene).

### 2.8. Immunolocalisation of rFhENO in whole *F. hepatica* NEJ and sections of adult worms

Immunolocalisation was carried out on fixed whole FhNEJ. Metacercariae (Ridgeway Research Ltd) were excysted and cultured for three hours as described by De Marco Verissimo et al. [34]. FhNEJ were fixed in 4% paraformaldehyde (PFA) for one hour at RT and then washed in AbD buffer (PBS containing 0.1% Triton X-100 (*v/v*), 0.1% bovine serum albumin (BSA, (*w/v*), and 0.1% NaN_3_ (*w/v*)). FhNEJ were probed overnight at 4°C in a 1:500 dilution of either of the following primary antibodies raised in rabbit: anti-rFhENO, pre-immune anti-rFhENO, anti-*F. hepatica* cathepsin L3 (rFhCL3; Eurogentec) as a non-related control, or pre-immune anti-rFhCL3. After three washes in AbD buffer, FhNEJ were probed with the secondary antibody, fluorescein isothiocyanate (FITC)-labelled goat-anti-rabbit IgG (1:200; Sigma-Aldrich), and incubated overnight at 4°C in the dark. The musculature of the FhNEJ was counterstained with phalloidin-tetramethyl rhodamine isothiocyanate (TRITC, Sigma-Aldrich) at a concentration of 200 µg/mL, overnight at 4°C in the dark. Stained FhNEJ were mounted onto slides using mounting solution (10% glycerol solution with 0.1 M propyl gallate) and images were taken on a Fluorview 3000 laser scanning confocal microscope (Olympus).

rFhENO was localised in the tissues of adult *F. hepatica* using immunofluorescence microscopy on JB-4 resin sections as described by Calvani et al. [35]. Briefly, adult parasites were fixed in 4% PFA for four hours at RT, washed in PBS, and dehydrated using increasing volumes of ethanol before being permeated with JB-4 resin (Sigma-Aldrich). Parasite sections were cut at 1 µm thickness, mounted on slides and probed with the following primary rabbit antibodies: anti-rFhENO (1:250 dilution) (David’s Biotechnologie GmbH), anti-*F. hepatica* cathepsin L1 pro-peptide (rFhCL1pp; 1:500 dilution, non-related control) and pre-immune anti-rFhCL1pp (1:500 dilution) (Eurogentec). Parasite sections were incubated at RT for five hours in a humid container. After three washes with PBST, the secondary antibody (goat anti-rabbit IgG FITC-conjugated diluted 1:200) was added and the samples were incubated overnight at 4°C in a dark, humid container. Following three further washes, sections were covered with mounting solution and a coverslip (thickness 0.13mm - 0.16mm, DeltaLab), and fluorescent labelling was visualized using the Leica DM2500 LED optical fluorescent microscope (Leica Microsystems).

### 2.9. Assessment of the immunogenicity of FhENO in sheep experimentally infected with *F. hepatica* by Western Blot

Sera were obtained from sheep experimentally infected with 120 *F. hepatica* metacercariae (Italian isolate; Ridgeway Research Ltd), previously described by López Corrales et al. [36]. The recombinant proteins, rFhENO, and rFhGAPDH2 as a related control, were resolved by SDS-PAGE (Bio-Rad) and electro-transferred onto a nitrocellulose membrane as described above. The membranes were blocked overnight at 4°C in blocking buffer and probed with pooled sheep sera obtained at time 0-, 7-, 11-, and 15-weeks post infection (WPI) (1:200 dilution) for one hour at RT. After five washes with PBST, the membranes were incubated with donkey anti-sheep IgG HRP-conjugated (1:15,000 dilution; Thermo Fisher Scientific) for one hour at RT. The blots were developed with SigmaFast 3,3’-Diamino-benzidine substrate (Sigma-Aldrich) and imaged using the G:BOX Chemi XRQ imager (Syngene).

### 2.10. Fibronectin, laminin and plasminogen binding assays

The ability of rFhENO to bind host FN, LM and/or PLG was assayed by enzyme-linked immunosorbent assays (ELISA) as described in Serrat et al. [26,27]. Briefly, multi-well microplates (Costar) were coated with 0.5 µg/well of rFhENO or 1% BSA as negative control, blocked and then incubated with increasing amounts of human FN (Santa Cruz Biotechnology) (from 0 µg to 2 µg), LM (Santa Cruz Biotechnology) (from 0 µg to 2 µg) or PLG (Origene) (from 0 µg to 3 µg). After incubation with the corresponding antibodies and chromogen [26,27], optical densities (OD) were measured at 492 nm in a Multiskan GO spectrophotometer (Thermo Fisher Scientific). A competition assay to study the involvement of lysine residues in PLG binding was performed by including 0-100 mM of the lysine analogue ε-aminocaproic acid (ε-ACA) during PLG incubation with rFhENO. The assays were performed in technical triplicates.

### 2.11. Plasminogen activation assays

rFhENO was tested in a PLG activation assay to explore the enhancement of plasmin generation by this recombinant protein following the methodology described by Serrat et al. [26]. Briefly, wells containing 2 µg of human PLG (Origene) were incubated with 3 µg of D-Val-Leu-Lys 4-nitroanilide dihydrochloride chromogenic substrate (S-2251) (Sigma-Aldrich) in the presence of 1 µg of rFhENO or 1% BSA as negative control in a test volume of 100 µL. Activation of PLG was initiated by the addition of 15 ng or 10 ng of t-PA (Sigma-Aldrich) or u-PA (Sigma-Aldrich), respectively, and plasmin generation was also measured in the absence of PLG activators. Plates were incubated at 37 °C for three hours and the amidolytic activity of generated plasmin was monitored by measuring the hydrolysis of the chromogenic substrate at 405 nm absorbance every 30 min in a Multiskan GO spectrophotometer (Thermo Fisher Scientific). Each sample was analysed in technical triplicate.

### 2.12. Statistical analysis

Plots were created with Prism 9 software (GraphPad Software) and statistical analyses were performed with the R Commander package [37]. Comparisons between two groups was performed with an unpaired Student’s *t*-test, and comparisons between three or more groups was performed with an Analysis of Variance (ANOVA) test followed by a Tukey post-hoc analysis for pair-wise comparisons. Unless otherwise stated, the differences were not significant.

## 3. RESULTS

### 3.1. Expression and purification of rFhENO

The FhENO gene was successfully expressed in ClearColi BL21 as an expected 47 kDa protein, with increasing expression overtime following induction with IPTG (Fig. 1A). Immunoblotting performed with anti-HisTag antibody detected the rFhENO protein successfully at the same MW, confirming its expression in the culture (Fig. 1B). We were able to successfully purify rFhENO (>95% purity) using NiNTA-Agarose affinity chromatography (Fig. 1C and D), which resulted in a yield of 12.8 mg/L of culture.

**Figure 1.**
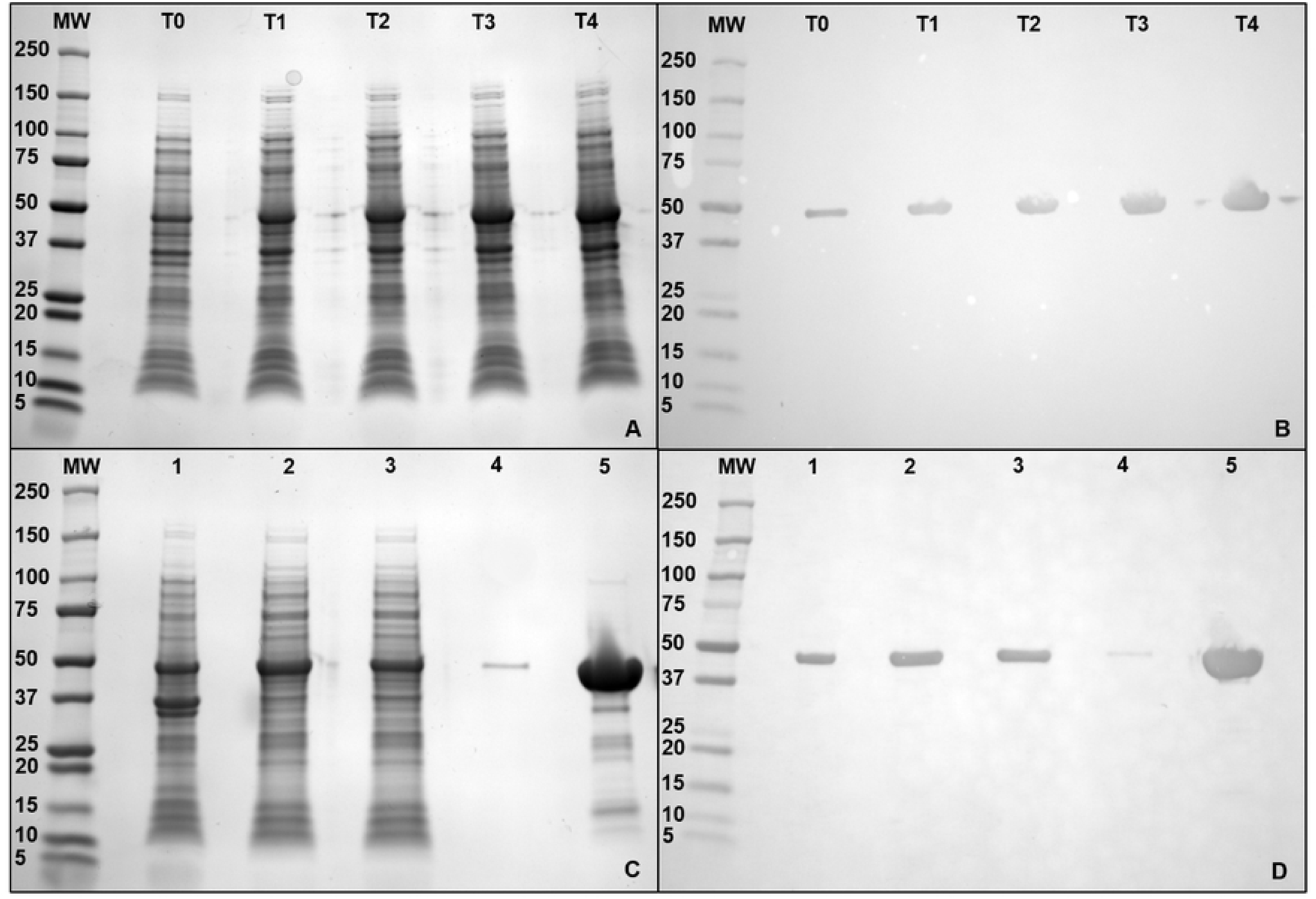
Recombinant expression and purification of *F. hepatica* enolase. **(A)** 4-20% SDS-PAGE gel of time points for recombinant expression of rFhENO in BL21 ClearColi system. T0, T1, T2, T3, and T4 refer to 0, 1, 2, 3, and 4 hours post induction with IPTG, respectively. **(B)** Corresponding immunoblot of time points of recombinant expression probed with an anti-polyhistidine antibody at 1:10,000 dilution. **(C)** 4-20% SDS-PAGE gel of NiNTA-Agarose affinity chromatography purification steps of rFhENO; (1) cell pellet of T4, (2) supernatant after cell lysis, (3) run-through of cell lysate, (4) wash of the purification column, and (5) rFhENO eluted from the column with 250 mM imidazole. **(D)** Corresponding immunoblot of affinity chromatography purification steps probed with an anti-polyhistidine antibody at 1:10,000 dilution. MW = molecular weight in kilodaltons.

### 3.2. rFhENO is functionally active as a homodimeric enzyme

SEC determined that rFhENO is primarily a dimer, with elution of 86.5% of the protein in the homodimeric form (peak 9.81), whilst 13.5% of it was observed in the monomeric form (Peak 12.16; Fig. 2). Specific activity assays showed that the enzymatic activity of rFhENO is associated with its dimeric structure, which can generate 8.07 nmole H_2_O_2_ / µM of dimeric rFhENO, while no hydrogen peroxide was generated by its monomeric form. For comparison, rFhENO, which did not undergo SEC separation, was also assayed for activity and generated 2.92 nmole H_2_O_2_ / µM rFhENO (Fig. 2, inset). The homodimeric form of rFhENO is 176.71% more active than the whole rFhENO purified by affinity chromatograph only, indicating that the enzyme was further purified by SEC.

**Figure 2.**
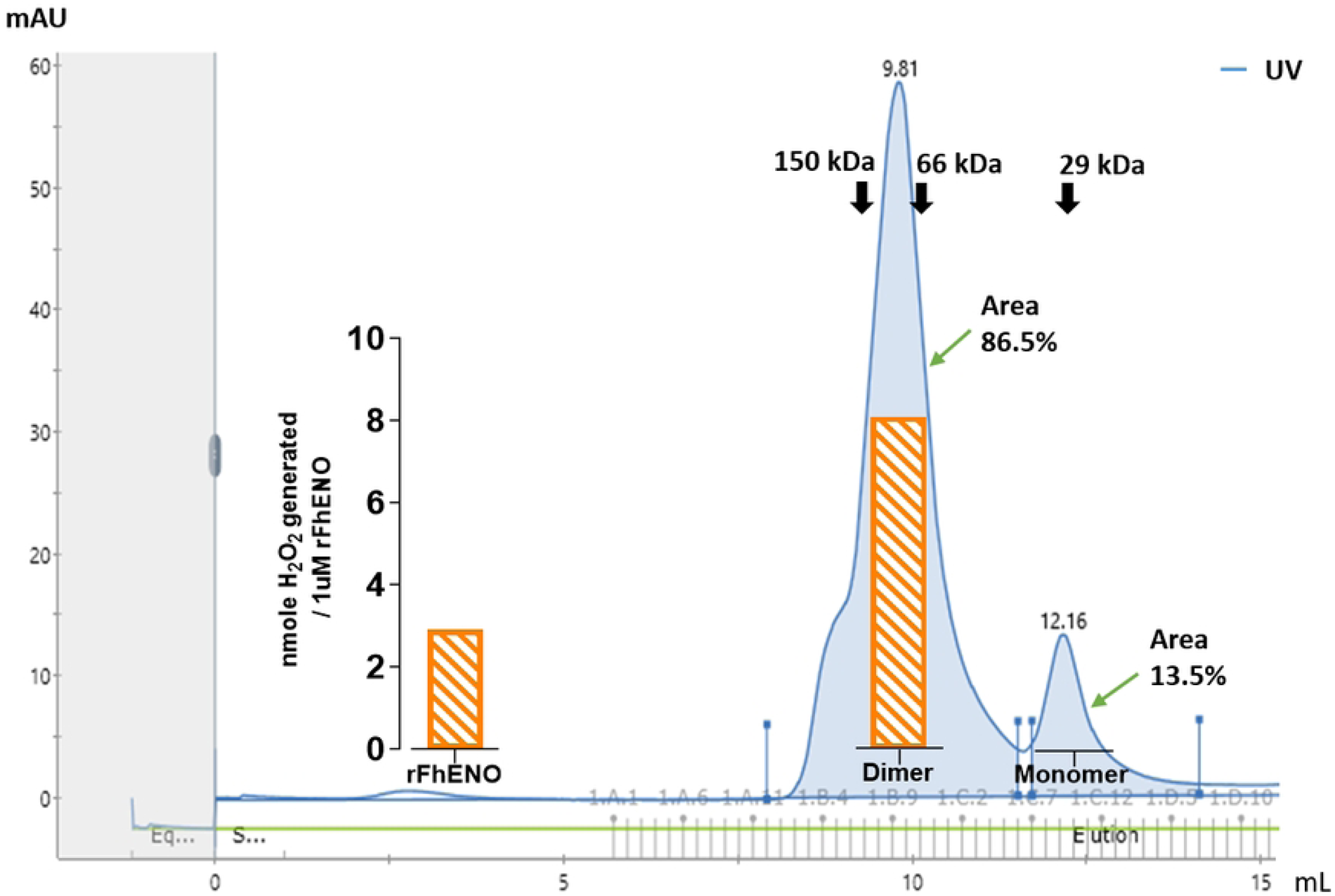
Size-exclusion chromatography (SEC) and enzyme activity assay of rFhENO. A SEC readout of separated dimeric and monomeric peaks of rFhENO are calculated to be ∼94 kDa and ∼47 kDa, respectively. The black arrows indicate the molecular weight size markers, alcohol dehydrogenase (150 kDa), bovine serum albumin (66 kDa), and carbonic anhydrase (29 kDa), resolved on the same SEC. Peak 9.81 (dimer) eluted at 9.81 mL between the molecular markers 150 and 66 kDa, indicating a dimeric form. Peak 12.16 (monomer) eluted at 12.16 mL between the molecular weight markers 66 and 29 kDa, indicative of the monomeric form. The fractions that eluted within these two peaks were pooled and assayed for enzyme activity at 0.1 µM. The rFhENO activity is graphically represented in orange and expressed as nmoles H_2_O_2_ generated per 1 µM of enzyme. Total purified rFhENO not separated by SEC was also assayed at 0.1 µM and its activity is expressed in the figure (inset on the left), to serve as a comparison of enzyme activity of the eluted peaks.

### 3.3. Enolase is present in the tegument and excretory/secretory extracts of adult *F. hepatica*

Western blot analysis was carried out on adult *F. hepatica* extracts, namely Soma, Teg, total ES, and concentrated ES fraction (fFhES), to infer the expression, localisation and function of FhENO within parasite tissues. Native FhENO was detected in the Soma and Teg extracts at 47 kDa, compatible with the molecular size of our recombinant enolase. An additional band at ∼35 kDa, possibly a breakdown product, was also detected by our polyclonal antibodies (Fig. 3A). Notably, FhENO was not detected in the total ES preparation; however, it was detected in the size-exclusion concentrated fraction of this extract, fFhES, with the same bands plus a few extra bands recognized in the Soma and Teg extracts (Fig. 3A). No bands were detected when the same extracts were probed with the pre-immune rabbit sera (Fig. 3B).

**Figure 3.**
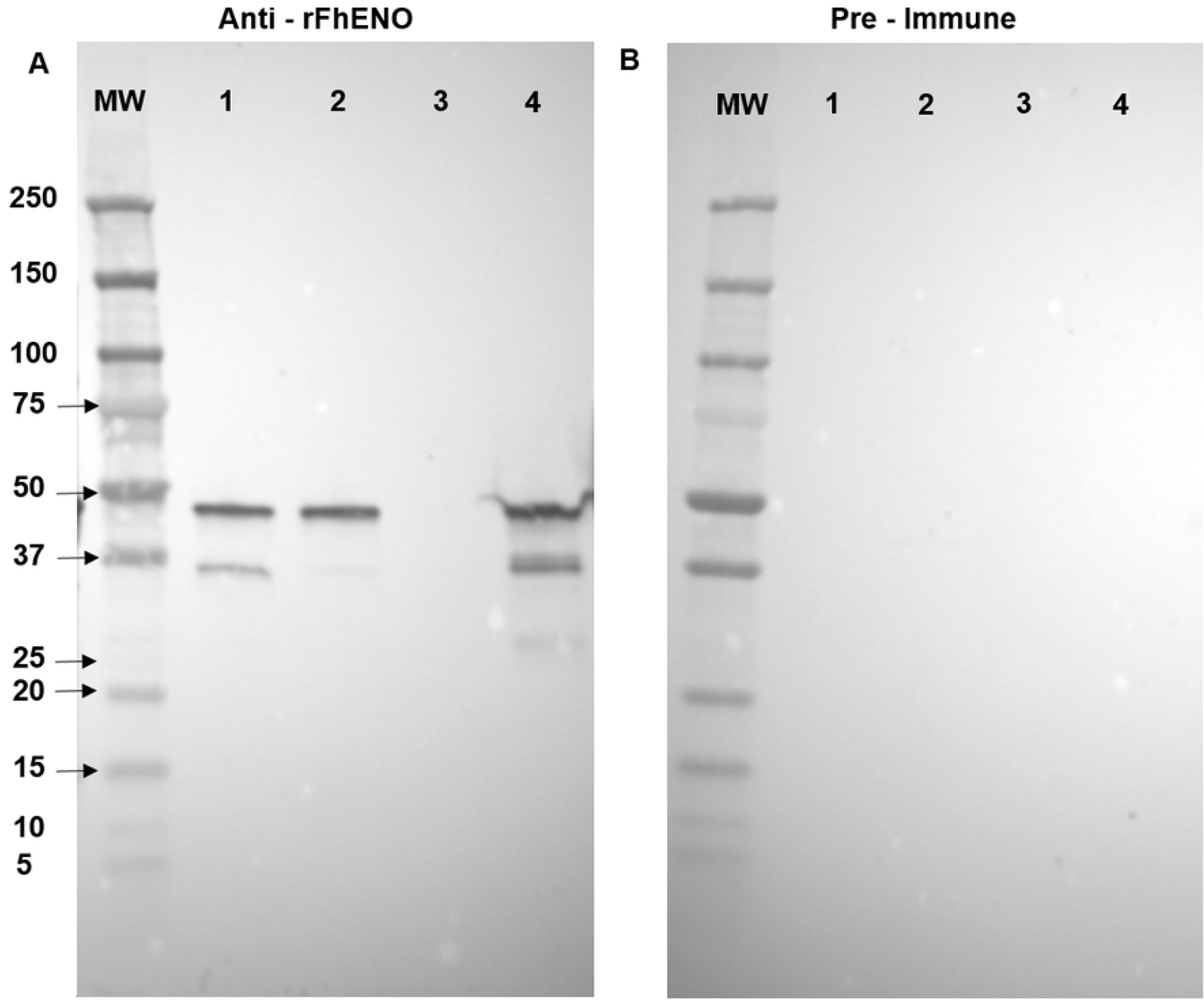
Detection of *F. hepatica* enolase in adult fluke extracts by immunoblot. Samples of the different adult liver fluke extracts (2 µg/lane) were probed with **(A)** Rabbit anti-rFhENO polyclonal antibody at a 1:1,000 dilution. **(B)** Pre–immune rabbit sera at a dilution of 1:1,000. MW: molecular weight in kilodaltons, (1) adult somatic extract (FhSoma), (2) adult tegumental extract (FhTeg), (3) total adult excretory/secretory products (FhES), and (4) concentrated size-separated fraction excretory secretory products (fFhES).

### 3.4. Immunolocalisation of FhENO in the tegument of *F. hepatica* NEJ and adult worms

Immunolocalisation studies carried out on FhNEJ using anti-rFhENO antibody revealed the presence of FhENO on the surface tegument (Fig. 4). FhNEJ probed with control anti-FhCL3 shows, as expected, the presence of FhCL3 in the gut of the parasite, whilst parasites probed with pre-immune sera showed no distinct labelling (Fig. 4). Immunolocalisation studies carried out on JB-4 resin-embedded sections of adult *F. hepatica*, showed a distinct presence of FhENO on the tegument layer of the parasite, mainly noticeable on the outer tegument of adults, covering even the spines as shown in the insets in Fig. 5D. In contrast, the positive control anti-FhCL1pp localised the major secreted protease in the gut of the parasite (Fig. 5B). Both sections probed with pre-immune sera showed only background fluorescence (Fig. 5A and C).

**Figure 4.**
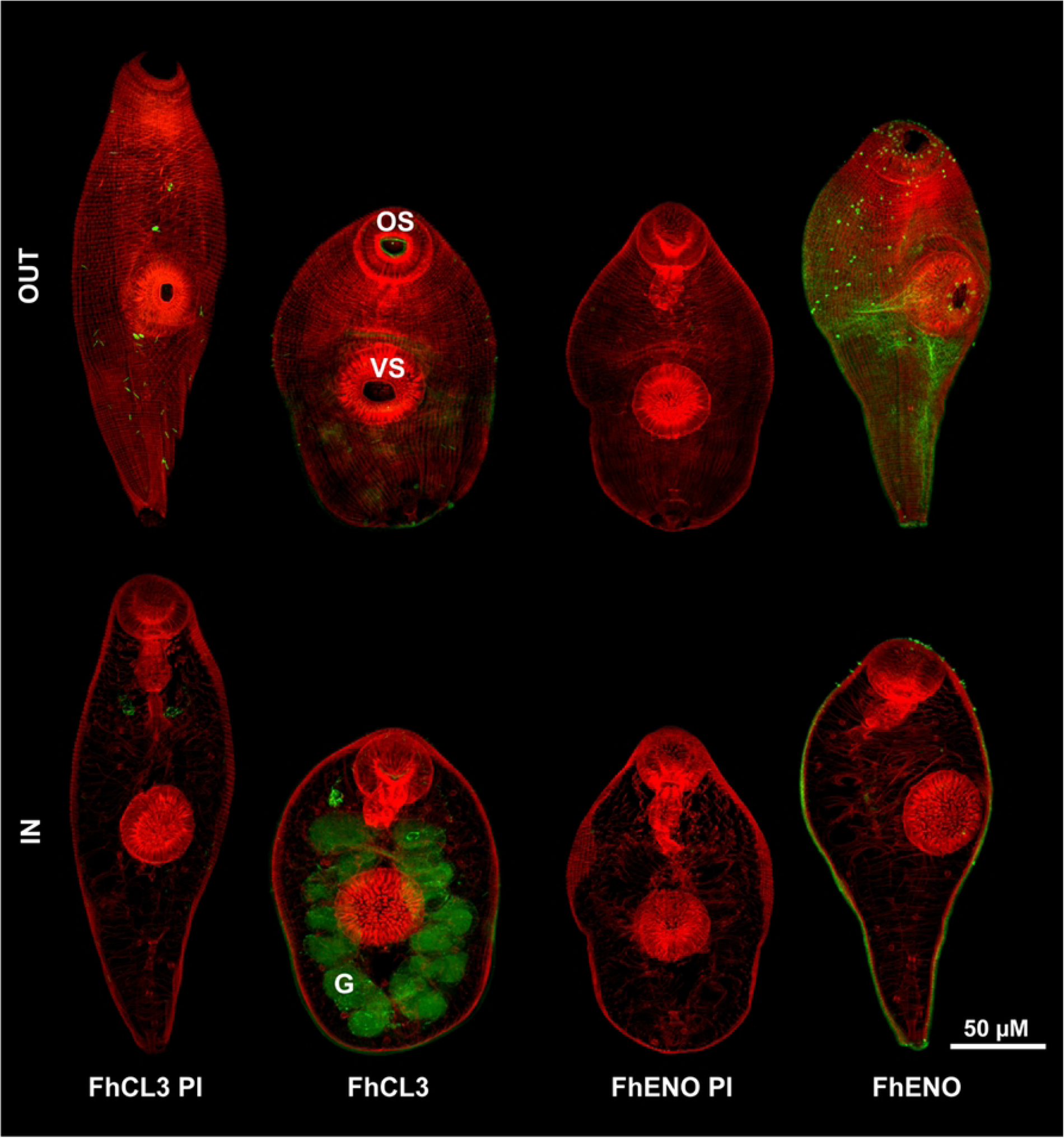
Immunolocalisation of native *F. hepatica* enolase in FhNEJ. Confocal microscopy of FhNEJ probed with pre-immune anti-rFhCL3 sera (FhCL3 PI), anti-rFhCL3 polyclonal antibody (FhCL3), pre-immune anti-rFhENO sera (FhENO PI), and anti-rFhENO polyclonal antibody (FhENO), from left to right. Positive antibody binding is show in green, whilst the musculature of the parasites was counter-stained with TRITC, which appears as red labelling. OUT and IN indicate the external and internal surfaces of the parasites, respectively. The main NEJ’s features are highlighted: G: Gut; OS: Oral sucker; VS: Ventral sucker. Scale bar: 50 μM.

**Figure 5.**
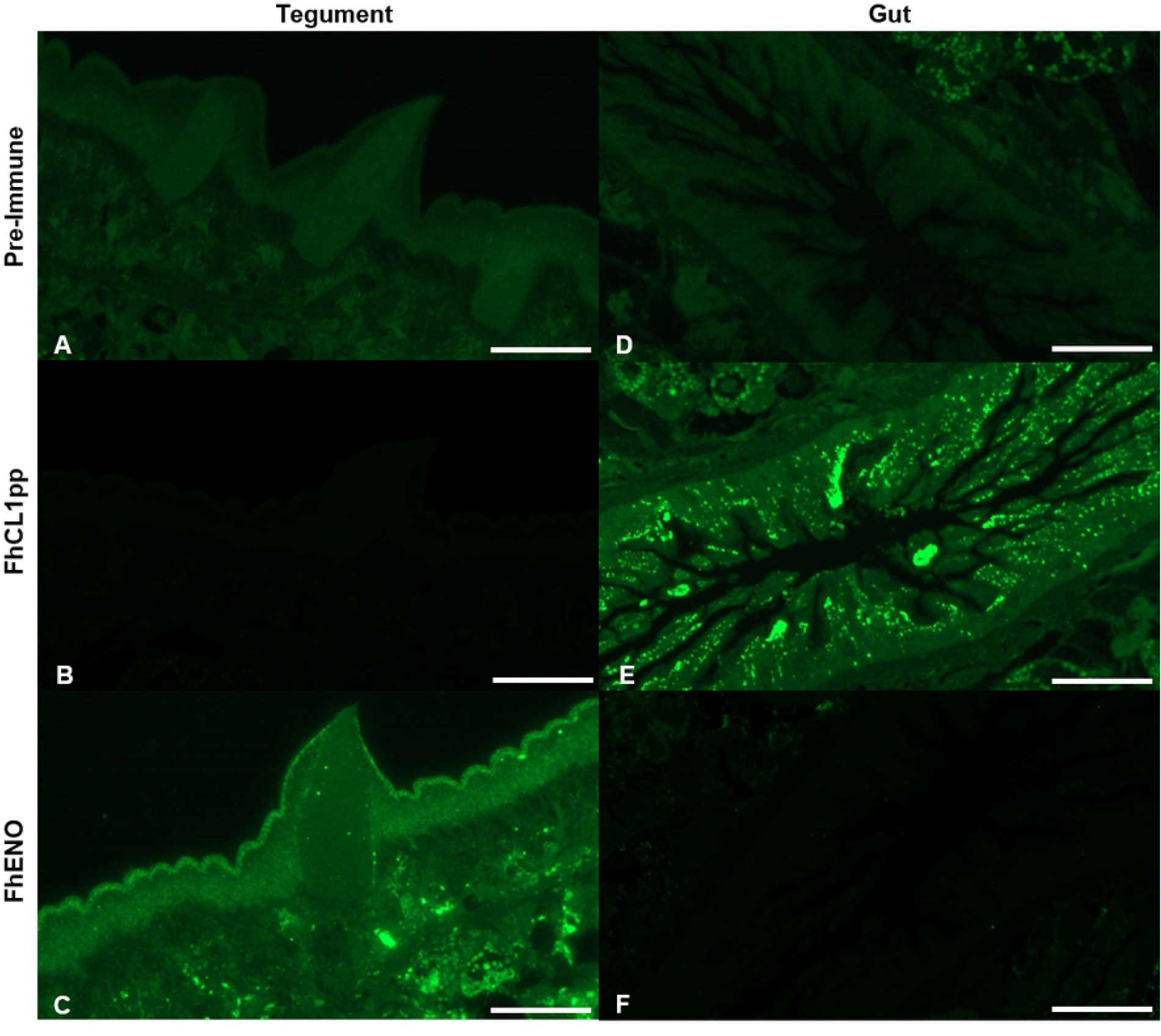
Immunolocalisation of native *F. hepatica* enolase in the tegument of adult *F. hepatica*. *F. hepatica* adult JB-4 resin-embedded sections in the area of the tegument (A – C) and gut (D – F). Sections were probed with **(A and D)** pre-immune anti-rFhCL1pp sera, **(B and E)** anti-rFhCL1pp polyclonal antibody, **(C and F)** anti-rFhENO polyclonal antibody. Scale bar: 50 μM.

### 3.5. *F. hepatica* enolase stimulates antibody response in sheep infected with *F. hepatica*

The antibody response elicited towards FhENO in sheep experimentally infected with *F. hepatica* was analysed by immunoblotting against rFhENO. We compared the antigenicity of rFhENO with another glycolytic enzyme, the GAPDH, which we have also recombinantly produced in our laboratory (rFhGAPDH). We found that under the conditions used, a background detection of both rFhENO and rFhGAPDH can be observed at 0 WPI. A definitive antibody response against rFhENO (strong band developed) was observed from 7 WPI, with an increase in the band recognition over the sampling period. A similar pattern was observed with rFhGAPDH; however, a much weaker band developed, suggesting that FhENO is more immunogenic in sheep during liver fluke infection.

### 3.6. rFhENO binds host laminin but not fibronectin

The adhesin property of the *F. hepatica* rFhENO was assessed by examining its ability to bind major components of the host ECM, such as FN and LM, using a protein-protein interaction assay (Fig. 7). Using this approach, we showed that rFhENO does not bind FN, which displayed comparable results to the negative controls, but, in contrast, the enzyme bound LM in concentration-dependent manner. Wells coated only with BSA, which served as negative controls, exhibited some non-specific binding activity, but the binding values were significantly lower (p >0.001) than those obtained by rFhENO.

**Figure 6.**
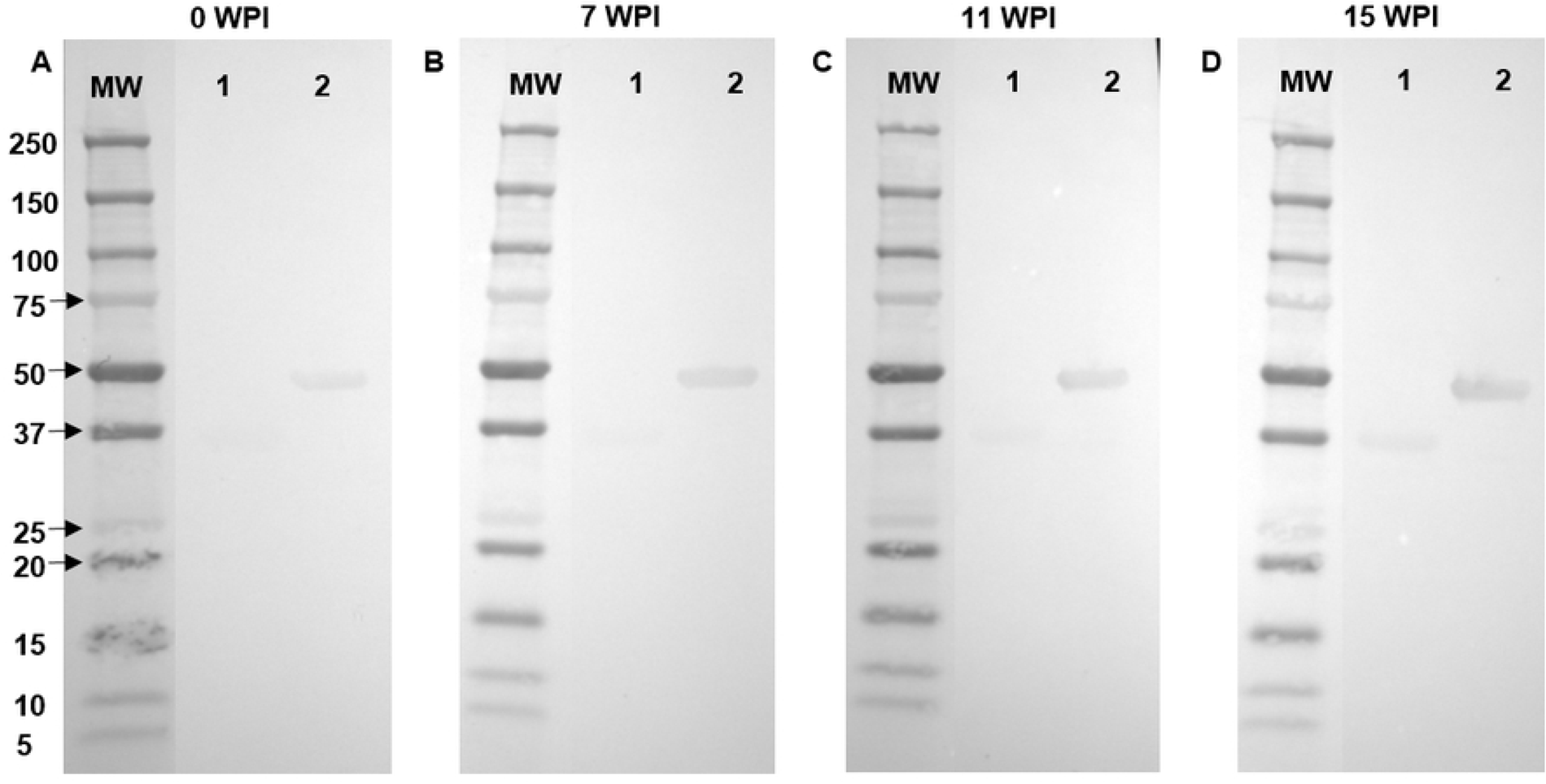
Immunogenicity of *F. hepatica* enolase in sheep experimentally infected with *F. hepatica*. Sera collected at different time points from sheep experimentally infected with *F. hepatica* was analysed for the presence of antibodies against rFhENO, as well as to another glycolytic enzyme, the glyceraldehyde-3-phosphate dehydrogenase (rFhGAPDH), for comparison. **(A)** 0 weeks post infection (WPI), **(B)** 7 WPI, **(C)** 11 WPI, and **(D)** 15 WPI. The recombinant antigens, rFhGAPDH (lane 1) and rFhENO (lane 2). MW: molecular weight in kilodaltons.

**Figure 7.**
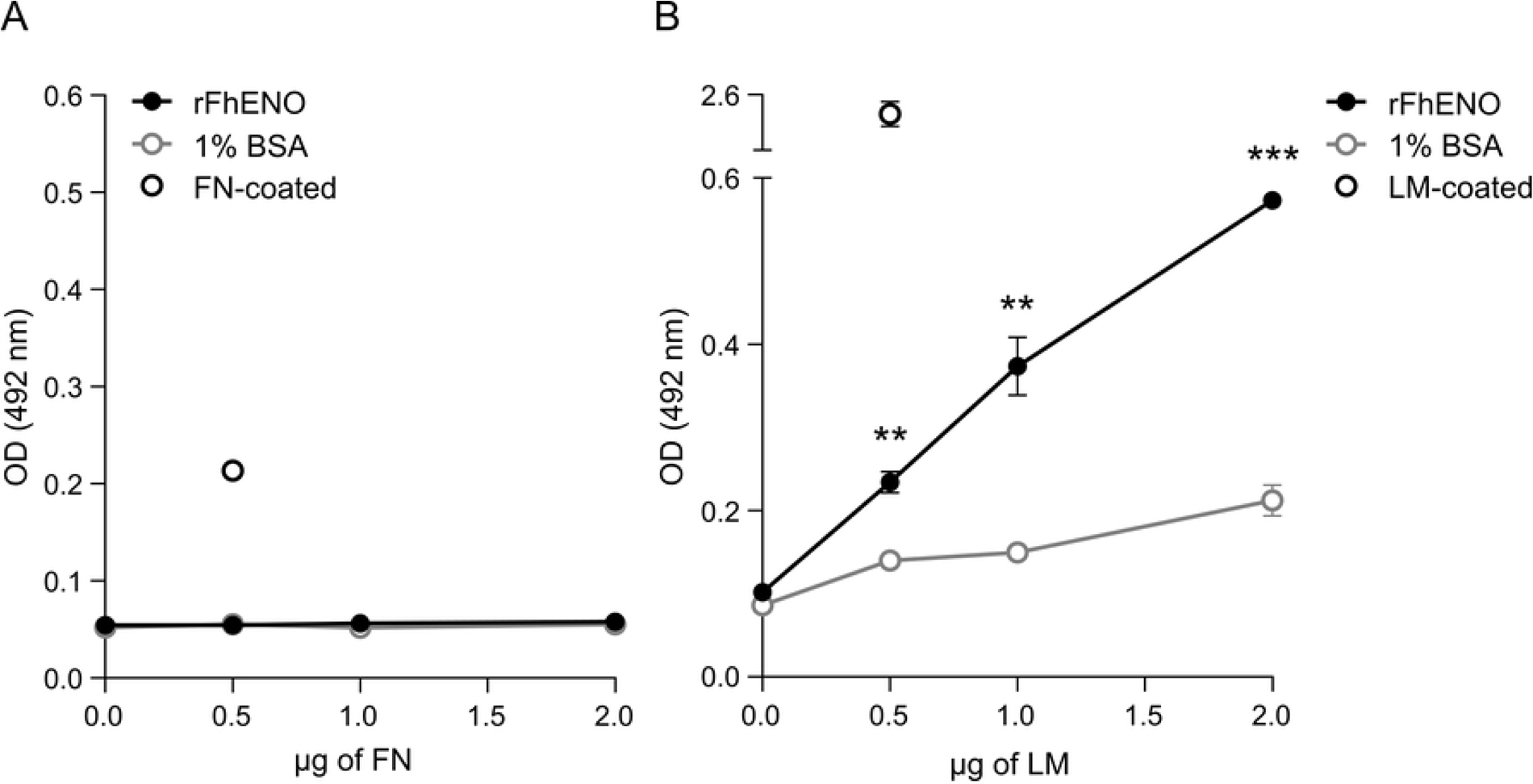
Protein-protein interaction assays to examine ability of rFhENO to bind fibronectin (FN) and laminin (LM). Binding of rFhENO to FN (A) and LM (B) was analysed over a range of FN and LM concentrations using a microtiter plate method. Wells coated with 1% BSA serve as negative controls, and wells coated with FN or LM serve as positive controls for antibody binding. Each data point represents the mean ± SD of three technical replicates. Asterisks indicate significant differences between rFhENO and 1% BSA (**p≤0.01, ***p≤0.001; Student’s *t*-test).

### 3.7. rFhENO binds host plasminogen and augments its activation to plasmin

Using a microtitre plate assay we showed that rFhENO binds PLG in a directly proportional manner to the amount of host PLG added to the plate well (Fig. 8A). To determine whether binding of rFhENO to PLG occurs via the PLG kringle domains, which interacts with lysine residues of partner proteins, a competition experiment was carried out in the presence of the lysine analogue ε-ACA (Fig. 8B). PLG binding to rFhENO was inhibited by ∼80% after the addition of ε-ACA, which completely abrogated PLG binding to rFhENO at 25 mM. The characteristics of rFhENO interaction with PLG were further assessed by measuring the amidolytic activity of plasmin generated by the interaction of these two factors in the presence or absence of the PLG activators t-PA and u-PA. As shown by the results, both PLG activators can generate active plasmin on their own, but this activity was significantly enhanced in the presence of rFhENO (Fig. 8C and D).

**Figure 8.**
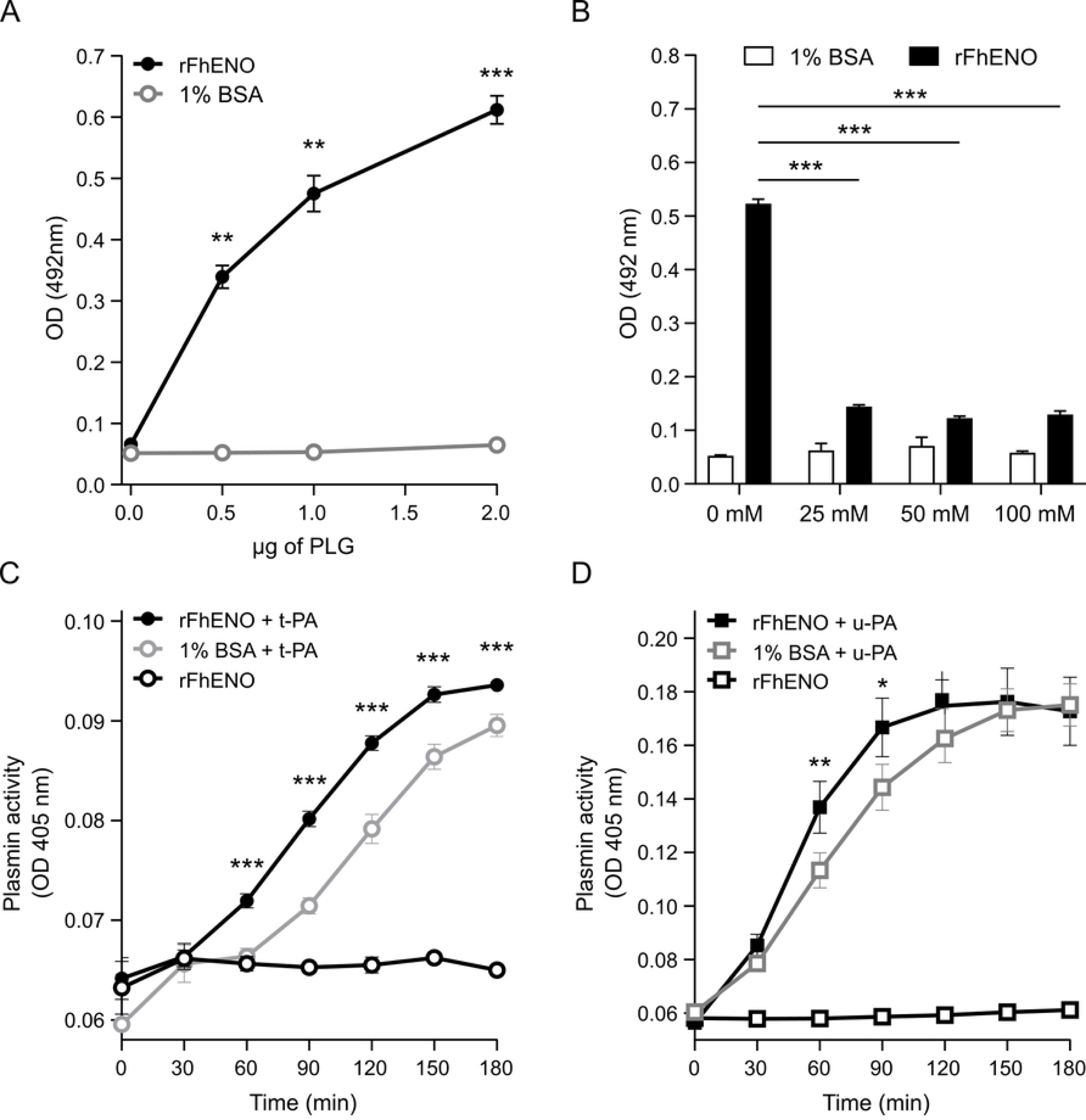
Plasminogen (PLG) binding to rFhENO and stimulation of plasmin generation from bound PLG. PLG binding to rFhENO was analysed using a microtiter plate method (A) in the presence or absence of the lysine analogue ε-ACA (B). rFhENO was incubated with PLG in the presence of t-PA (C) or u-PA (D) and a plasmin-specific chromogenic substrate and plasmin generation was measured by monitoring substrate cleavage over time. Wells containing 1% BSA instead of rFhENO serve as negative controls. Data points represent the mean ± SD of three technical replicates. Asterisks indicate significant differences between rFhENO and 1% BSA (*p≤0.05, **p≤0.01, ***p≤0.001; Student’s *t*-test [A], one-way ANOVA [B-D]).

## 4. Discussion

Moonlighting proteins are characterized by their multifunctionality, enabling them to participate in numerous cellular processes of high biochemical or biophysical relevance under the guise of the same polypeptide chain. These proteins are expressed across the evolutionary tree, performing diverse functions in response to environmental changes or different locations. Notably, intracellular enzymes integral to glycolysis (such as enolase) and other conserved metabolic pathways exhibit alternative functions when exposed on the cell surface. This phenomenon is recurrent in pathogenic organisms, where surface-exposed multifunctional proteins facilitate interactions with host tissues and molecules, vital for infection or virulence processes [38,39]. In this study, we aimed to characterize the moonlighting functions of the glycolytic protein enolase from the zoonotic parasitic trematode *F. hepatica*.

First, after confirming the nucleotide sequence of *F. hepatica* enolase, we obtained a peptide sequence of 47 kDa through the recombinant expression and downstream purification of rFhENO, with a MW similar to that obtained in the production of enolases in other helminth parasites such as *Schistosoma mansoni* (47 kDa) or *Trichinella spiralis* (51.95 kDa) [40,41]. Next, the characterization of rFhENO began with testing its canonical function as a cytosolic protein involved in the glycolytic pathway. This protein plays a critical role facilitating the conversion of 2PGA to PEP during the final stages of the glycolytic pathway. Enolase, functioning enzymatically, typically exists as a dimer (either homo-or heterodimers), comprised of two subunits positioned in an antiparallel orientation, facing each other [14]. In our study, we confirmed that rFhENO primarily exists as a homodimer, with 86.5% of the protein eluting in dimeric form via SEC. Additionally, specific activity assays demonstrated that the enzymatic activity of rFhENO is associated with its dimeric structure. Enolase activity was evaluated using a coupled enzyme assay, where 2PGA is converted to PEP, leading to the generation of an intermediate that reacts with a peroxidase substrate. This reaction yields a fluorometric product directly proportional to the detected enolase activity, thus validating the canonical activity of rFhENO.

Secondly, to determine if rFhENO has moonlighting functions with relevance “in vivo”, it is essential to establish whether this protein is expressed and/or localised in parasite tissues that closely interact with the host environment [42]. For that reason, we produced polyclonal antibodies against rFhENO and used them to test the expression of native enolase in different extracts of adult worms of *F. hepatica* by Western blot, as well as its localisation in the tissues of juvenile and adult parasites by microscopic techniques. Our assays revealed the expression of enolase in the total somatic extract of *F. hepatica*, and more specifically in extracts from the host-parasite interface, such as the tegumental and excretory/secretory extracts. These results are consistent with those obtained by other authors who identified enolase in somatic [43], tegumental [44], and excretory/secretory [45] extracts of adult *F. hepatica* worms using proteomic techniques. These expression levels were corroborated and expanded upon in immunolocalisation assays, where we observed how enolase is particularly present in the tegumental layer of both NEJ and adult worms of *F. hepatica*. This suggests the importance of this protein for *F. hepatica* in its interaction with its host throughout its life cycle, since despite the continuous turnover that the parasite tegument shows during parasite migration from its excystment in the intestine until it reaches the bile ducts [46], enolase maintains its expression at the host-parasite interface. The tegument of *F. hepatica* is recognized as dynamic, interacting continuously with host tissues to ensure survival of the parasite [21]. Composed of syncytial secretory cells, the *F. hepatica* tegument, including that of FhNEJ, undergoes significant changes in structure and composition during parasite maturation. Its secretions, crucial for migration and immune evasion, are released via conventional and unconventional pathways, such as exosome-mediated routes, at the host-parasite interface [47,48]. Indeed, enolase has been identified through proteomic techniques as part of the cargo of extracellular vesicles in both juvenile [49] and adult worms [50] of *F. hepatica*, reinforcing the potential importance of this protein in the host-parasite interaction in fasciolosis.

After confirming the presence of *F. hepatica* enolase in the extracts of the host-parasite interface, we investigated whether this antigen could be recognized by the host’s immune system in an experimental infection. Western blot results using sera from sheep experimentally infected with the parasite showed that rFhENO is recognized from 7 WPI, revealing the immunogenicity of this protein before the parasites reach their final destination, the bile ducts. In the life cycle of *F. hepatica*, eggs released by mature worms are found in bile from 8 WPI, and later in faeces (9-12 WPI) [22], allowing direct diagnosis of fasciolosis through coprological techniques [51]. Therefore, early recognition of enolase by host antibodies could be explored for its potential to establish early serodiagnosis for fasciolosis. In this regard, enolase was identified by Morales and Espino [52] as one of the most immunoreactive antigens from a tegumental extract of *F. hepatica*, enabling serodiagnosis of human fasciolosis with high sensitivity, specificity, and accuracy. More recently, a recombinant enolase from *F. hepatica* has been tested for the serodiagnosis of fasciolosis in sheep by ELISA, yielding data with sensitivity and specificity rates of 90% and 97.14% [53]. However, these authors found that the recombinant form of enolase showed significant cross-reactions with other helminths in both Western blot and ELISA assays.

Enolase is an enzyme that exhibits a high level of sequence conservation across species, assuming it is present in all organisms, and due to the significance of the process in which it participates, it has not undergone profound changes [54]. This high degree of homology could hinder its use as a vaccine antigen. However, contrary to this assumption, Mahana et al. [55] demonstrated that a tegumental fraction of *F. gigantica*, consisting essentially of *F. gigantica* enolase, induced antibodies capable of binding to the surface membrane of NEJ and mediating their attrition, generating a significant protection of approximately 40% in a vaccine experiment conducted in sheep. This protein also showed 96-100% homology with *F. hepatica* enolase. These data, along with the fact that moonlighting proteins are promising anti-parasite candidate vaccines as they could interfere with a plethora of pathogen physiological functions, should not rule out rFhENO as a potential vaccine antigen against *F. hepatica*. Indeed, isoforms of this protein have been tested in various protection assays against other helminths such as *Ascaris suum*, *Clonorchis sinensis*, *Taenia pisiformis*, or *T. spiralis* with promising results [56–59].

The multifunctionality of enolase has been known for over 35 years [60], and its role as a moonlighting protein in pathogenic organisms, especially in bacteria, with important functions related to the invasion process, has been extensively studied [61]. In this sense, enolase has been reported to bind a significant amount of host proteins, especially ECM proteins and PLG, justifying the enzyme’s characterisation as an adhesin that is part of bacterial interactions with the environment and responses to environmental changes [16–18,20]. Although the ability of enolase to bind PLG and interact with the host fibrinolytic system has been reported in some helminth species in recent years [19], its role as an adhesin against host ECM proteins has not yet been explored in this group of parasites.

Our results show that rFhENO has the ability to bind LM, but not FN, in a concentration-dependent manner. This could be significant at the intestinal level since LM constitutes a significant part of the intestinal basement membrane, located directly beneath the intestinal epithelium, while FN is predominantly present in deeper tissue layers, such as the interstitial matrix [5]. These findings suggest that *F. hepatica* demonstrates a specific adaptation to interact with the tissue microenvironment encountered immediately after excystment. Furthermore, this hypothesis could be extrapolated to the liver since this pattern of organization of the ECM is also maintained at the hepatic level [62], the tissue through which *F. hepatica* migrates several weeks until the parasite enters the bile ducts to mature into the adult stage [22]. These results also agree with those recently obtained by our group in which we observed the ability of an extract from FhNEJ tegument to bind LM but not FN as a potential invasion mechanism at the intestinal level [27].

Our data also demonstrate the ability of rFhENO to interact with the host fibrinolytic system by binding to PLG and enhancing its conversion to plasmin in the presence of the activators t-PA and u-PA. Competition assays using ε-ACA acid indicated the participation of lysine residues from rFhENO in the binding of PLG. The association between PLG and its receptors has been linked to the presence of carboxyl-terminally lysine residues [63], so the mechanism of binding of the parasitic protein to PLG is similar to what occurs physiologically. Plasmin, the end product of the fibrinolytic system, exhibits broad and potent proteolytic activity and degrades both fibrin and various constituents of the ECM and connective tissue. Consequently, the enlistment of this enzyme by any blood or tissue pathogen may not only serve as an effective mechanism to evade potential immobilization by fibrin blood clots but also assist the pathogen in its dissemination and establishment within the host through the breakdown of diverse ECM components [9]. In this context, a large number of blood and tissue parasites express PLG receptors on their surface or secrete them, allowing them to interact with the fibrinolytic system of their hosts. In fact, enolase has been recently reported as the most commonly identified PLG receptor in parasitic helminths in a scoping review conducted by our group [19]. Our results also confirm those obtained twenty years ago by Bernal et al. [64], who identified an enolase in the excretory/secretory products of *F. hepatica* as a potential PLG receptor, and complement our own findings in which we characterize the potential of the tegument of *F. hepatica* juvenile [26] and adult worms [65] to interact with the fibrinolytic system of their host. Indeed, the utilization of this mechanism throughout the life cycle of *F. hepatica* could have different functions adapted to the changing environments faced by the parasite. Enhancing the proteolytic capacity of plasmin could be related to the invasive capacity of the juvenile worm of *F. hepatica* due to the ability of this host protease to degrade ECM proteins. However, it could also represent a strategy associated with the nutritional requirements of the adult worm, an obligate blood feeder, for which activation of the host’s fibrinolytic system would allow it to dissolve blood clots of fibrin, as described for other helminth parasites [9,19]. Furthermore, the enhancement of plasmin generation by rFhENO occurs in the presence of both activators u-PA and t-PA, which primarily operate at tissue and vascular levels [66], respectively, reinforcing the hypothesis of the key role of this mechanism for *F. hepatica* in both its migration and nutrition.

In conclusion, we have produced a functional recombinant form of *F. hepatica* enolase and characterized its role as a moonlighting protein. rFhENO is active as a homodimeric glycolytic enzyme fulfilling its canonical function, but it also functions as an adhesin capable of binding host ECM proteins and acts as a receptor for PLG, enhancing the activity of the broad-spectrum protease, host plasmin. This, together with its demonstrated expression at the host-parasite interface in both the juvenile and adult stages of the parasite, reinforces its potential involvement in disparate mechanisms such as the migration or nutrition of *F. hepatica*. The involvement of enolase in processes crucial for the survival of *F. hepatica*, coupled with its demonstrated immunogenicity even in early stages, strengthens the possibility of further investigation as a diagnostic marker or as a therapeutic target or vaccine antigen with the ultimate aim to establish effective management and control measures against both animal and human fasciolosis.

## Acknowledgements

JGM acknowledges funding received from projects ULYSSES (RTI2018-093463-J-100) and URANUS (CNS2022-135561) funded by the Spanish Ministry of Science and Innovation. IRNASA-CSIC group acknowledges funding received from Project “CLU-2019-05 - IRNASA-CSIC Unit of Excellence”, funded by the Junta de Castilla y León and cofounded by the European Union (FEDER “Europe drives our growth”) and funding from the Programme for strengthening research structures “Stairway to excellence” internationalisation aid, cofounded by the Junta de Castilla y León and the European Regional Development Fund. JS acknowledges the support of the Junta de Castilla y León for her Predoctoral contract. KC, NEDC, HJ, AF, RL, EOK, and JPD were supported by a Science Foundation Ireland Professorial grant, 17/RP/5368 and CDMV by the Irish Research Council, RCS1904. The authors acknowledge the facilities and scientific and technical assistance of the Centre for Microscopy & Imaging at the University of Galway (https://imaging.universityofgalway.ie/imaging/).

## References

1. Read AF, Skorping A. The evolution of tissue migration by parasitic nematode larvae. Parasitology. 1995 Sep;111 (Pt 3): 359–371. doi: 10.1017/s0031182000081919.

2. Mulcahy G, O’Neill S, Fanning J, McCarthy E, Sekiya M. Tissue migration by parasitic helminths - an immunoevasive strategy? Trends Parasitol. 2005 Jun;21(6): 273–277. doi: 10.1016/j.pt.2005.04.003.

3. 3. Rohde M, Cleary PP. Adhesion and invasion of Streptococcus pyogenes into host cells and clinical relevance of intracellular streptococci. In: Ferretti JJ, Stevens DL, Fischetti VA, editors. Streptococcus pyogenes: Basic Biology to Clinical Manifestations. Oklahoma City (OK): University of Oklahoma Health Sciences Center; 2022 Oct 8. Chapter 17.

4. Vaca DJ, Thibau A, Schütz M, Kraiczy P, Happonen L, Malmström J, et al. Interaction with the host: the role of fibronectin and extracellular matrix proteins in the adhesion of Gram-negative bacteria. Med Microbiol Immunol. 2020 Jun;209(3): 277–299. doi: 10.1007/s00430-019-00644-3.

5. Pompili S, Latella G, Gaudio E, Sferra R, Vetuschi A. The Charming World of the Extracellular Matrix: A Dynamic and Protective Network of the Intestinal Wall. Front Med (Lausanne). 2021 Apr 16;8: 610189. doi: 10.3389/fmed.2021.610189.

6. Kasný M, Mikes L, Hampl V, Dvorák J, Caffrey CR, Dalton JP, et al. Chapter 4. Peptidases of trematodes. Adv Parasitol. 2009;69: 205–297. doi: 10.1016/S0065-308X(09)69004-7.

7. Yang Y, Wen Yj, Cai YN, Vallée I, Boireau P, Liu MY, et al. Serine proteases of parasitic helminths. Korean J Parasitol. 2015 Feb;53(1): 1–11. doi: 10.3347/kjp.2015.53.1.1.

8. Grote A, Caffrey CR, Rebello KM, Smith D, Dalton JP, Lustigman S. Cysteine proteases during larval migration and development of helminths in their final host. PLoS Negl Trop Dis. 2018 Aug 23;12(8): e0005919. doi: 10.1371/journal.pntd.0005919.

9. González-Miguel J, Siles-Lucas M, Kartashev V, Morchón R, Simón F. Plasmin in Parasitic Chronic Infections: Friend or Foe? Trends Parasitol. 2016 Apr;32(4): 325–335. doi: 10.1016/j.pt.2015.12.012.

10. Cesarman-Maus G, Hajjar KA. Molecular mechanisms of fibrinolysis. Br J Haematol. 2005 May;129(3): 307–321. doi: 10.1111/j.1365-2141.2005.05444.x.

11. Pala ZR, Ernest M, Sweeney B, Jeong YJ, Pascini TV, Alves E Silva TL, et al. Beyond cuts and scrapes: plasmin in malaria and other vector-borne diseases. Trends Parasitol. 2022 Feb;38(2): 147–159. doi: 10.1016/j.pt.2021.09.008.

12. Gómez-Arreaza A, Acosta H, Quiñones W, Concepción JL, Michels PA, Avilán L. Extracellular functions of glycolytic enzymes of parasites: unpredicted use of ancient proteins. Mol Biochem Parasitol. 2014 Feb;193(2): 75–81. doi: 10.1016/j.molbiopara.2014.02.005.

13. Pancholi V. Multifunctional alpha-enolase: its role in diseases. Cell Mol Life Sci. 2001 Jun;58(7): 902–920. doi: 10.1007/pl00000910.

14. Díaz-Ramos A, Roig-Borrellas A, García-Melero A, López-Alemany R. α-Enolase, a multifunctional protein: its role on pathophysiological situations. J Biomed Biotechnol. 2012;2012: 156795. doi: 10.1155/2012/156795.

15. Avilán L, Gualdrón-López M, Quiñones W, González-González L, Hannaert V, Michels PA, et al. Enolase: a key player in the metabolism and a probable virulence factor of trypanosomatid parasites-perspectives for its use as a therapeutic target. Enzyme Res. 2011;2011: 932549. doi: 10.4061/2011/932549.

16. Li Q, Liu H, Du D, Yu Y, Ma C, Jiao F, et al. Identification of Novel Laminin- and Fibronectin-binding Proteins by Far-Western Blot: Capturing the Adhesins of *Streptococcus suis* Type 2. Front Cell Infect Microbiol. 2015 Nov 16;5: 82. doi: 10.3389/fcimb.2015.00082.

17. Salzillo M, Vastano V, Capri U, Muscariello L, Sacco M, Marasco R. Identification and characterization of enolase as a collagen-binding protein in *Lactobacillus plantarum*. J Basic Microbiol. 2015 Jul;55(7): 890–897. doi: 10.1002/jobm.201400942.

18. Wang J, Yu Y, Li Y, Li S, Wang L, Wei Y, et al. A multifunctional enolase mediates cytoadhesion and interaction with host plasminogen and fibronectin in *Mycoplasma hyorhinis*. Vet Res. 2022 Mar 25;53(1): 26. doi: 10.1186/s13567-022-01041-0.

19. Diosdado A, Simón F, Serrat J, González-Miguel J. Interaction of helminth parasites with the haemostatic system of their vertebrate hosts: a scoping review. Parasite. 2022;29: 35. doi: 10.1051/parasite/2022034.

20. Xie Q, Xing H, Wen X, Liu B, Wei Y, Yu Y, et al. Identification of the multiple roles of enolase as an plasminogen receptor and adhesin in *Mycoplasma hyopneumoniae*. Microb Pathog. 2023 Jan;174: 105934. doi: 10.1016/j.micpath.2022.105934.

21. González-Miguel J, Becerro-Recio D, Siles-Lucas M. Insights into *Fasciola hepatica* Juveniles: Crossing the Fasciolosis Rubicon. Trends Parasitol. 2021 Jan;37(1): 35–47. doi: 10.1016/j.pt.2020.09.007.

22. Andrews SJ, Cwiklinski K, Dalton JP. The discovery of Fasciola hepatica and its life cycle. In: Dalton JP, editor. Fasciolosis. Wallingford, UK: CAB International; 2022. pp. 18–46.

23. World Health Organization (2007). Report of the WHO Informal Meeting on use of Triclabendazole in Fascioliasis Control. World Health Organization, Geneva, Switzerland. WHO/CDS/NTD/PCT/2007.1.

24. Alvarez Rojas CA, Jex AR, Gasser RB, Scheerlinck JP. Techniques for the diagnosis of *Fasciola* infections in animals: room for improvement. Adv Parasitol. 2014;85: 65–107. doi: 10.1016/B978-0-12-800182-0.00002-7.

25. Molina-Hernández V, Mulcahy G, Pérez J, Martínez-Moreno Á, Donnelly S, O’Neill SM, et al. *Fasciola hepatica* vaccine: we may not be there yet but we’re on the right road. Vet Parasitol. 2015 Feb 28;208(1-2): 101–111. doi: 10.1016/j.vetpar.2015.01.004.

26. Serrat J, Becerro-Recio D, Torres-Valle M, Simón F, Valero MA, Bargues MD, et al. *Fasciola hepatica* juveniles interact with the host fibrinolytic system as a potential early-stage invasion mechanism. PLoS Negl Trop Dis. 2023 Apr 21;17(4): e0010936. doi: 10.1371/journal.pntd.0010936.

27. Serrat J, Torres-Valle M, López-García M, Becerro-Recio D, Siles-Lucas M, González-Miguel J. Molecular Characterization of the Interplay between *Fasciola hepatica* Juveniles and Laminin as a Mechanism to Adhere to and Break through the Host Intestinal Wall. Int J Mol Sci. 2023 May 3;24(9): 8165. doi: 10.3390/ijms24098165.

28. Davis RE, Singh H, Botka C, Hardwick C, Ashraf el Meanawy M, Villanueva J. RNA trans-splicing in Fasciola hepatica. Identification of a spliced leader (SL) RNA and SL sequences on mRNAs. J Biol Chem. 1994 Aug 5;269(31): 20026–20030.

29. Cwiklinski K, Dalton JP, Dufresne PJ, La Course J, Williams DJ, Hodgkinson J, et al. The *Fasciola hepatica* genome: gene duplication and polymorphism reveals adaptation to the host environment and the capacity for rapid evolution. Genome Biol. 2015 Apr 3;16(1): 71. doi: 10.1186/s13059-015-0632-2. P

30. Tran N, Ricafrente A, To J, Lund M, Marques TM, Gama-Carvalho M, et al. *Fasciola hepatica* hijacks host macrophage miRNA machinery to modulate early innate immune responses. Sci Rep. 2021 Mar 24;11(1): 6712. doi: 10.1038/s41598-021-86125-1.

31. Madeira F, Pearce M, Tivey ARN, Basutkar P, Lee J, Edbali O, et al. Search and sequence analysis tools services from EMBL-EBI in 2022. Nucleic Acids Research. 2022 Apr 12;50(W1). doi: 10.1093/nar/gkac240.

32. Cwiklinski K, Drysdale O, López Corrales J, Corripio-Miyar Y, De Marco Verissimo C, Jewhurst H, et al. Targeting secreted protease/anti-protease balance as a vaccine strategy against the helminth *Fasciola hepatica*. Vaccines (Basel). 2022;10(2): 155. doi: 10.3390/vaccines10020155.

33. Murphy A, Cwiklinski K, Lalor R, O’Connell B, Robinson MW, Gerlach J, et al. *Fasciola hepatica* Extracellular Vesicles isolated from excretory-secretory products using a gravity flow method modulate dendritic cell phenotype and activity. PLoS Negl Trop Dis. 2020 Sep 8;14(9): e0008626. doi: 10.1371/journal.pntd.0008626.

34. De Marco Verissimo C, Jewhurst HL, Tikhonova IG, Urbanus RT, Maule AG, Dalton JP, et al. *Fasciola hepatica* serine protease inhibitor family (serpins): Purposely crafted for regulating host proteases. PLoS Negl Trop Dis. 2020 Aug 6;14(8): e0008510. doi: 10.1371/journal.pntd.0008510.

35. Calvani NED, De Marco Verissimo C, Jewhurst HL, Cwiklinski K, Flaus A, Dalton JP. Two Distinct Superoxidase Dismutases (SOD) Secreted by the Helminth Parasite *Fasciola hepatica* Play Roles in Defence against Metabolic and Host Immune Cell-Derived Reactive Oxygen Species (ROS) during Growth and Development. Antioxidants. 2022 Oct 1;11(10): 1968. doi: 10.3390/antiox11101968.

36. López Corrales J, Cwiklinski K, De Marco Verissimo C, Dorey A, Lalor R, Jewhurst H, et al. Diagnosis of sheep fasciolosis caused by Fasciola hepatica using cathepsin L enzyme-linked immunosorbent assays (ELISA). Vet Parasitol. 2021 Oct 1;298: 109517. doi: 10.1016/j.vetpar.2021.109517.

37. Fox J, Marquez MM, Bouchet-Valat M. Rcmdr: R Commander. R package version 2.9–2. 2024. https://socialsciences.mcmaster.ca/jfox/Misc/Rcmdr/.

38. Jeffery CJ. Enzymes, pseudoenzymes, and moonlighting proteins: diversity of function in protein superfamilies. FEBS J. 2020 Oct;287(19): 4141–4149. doi: 10.1111/febs.15446.

39. Franco-Serrano L, Sánchez-Redondo D, Nájar-García A, Hernández S, Amela I, Perez-Pons JA, et al. Pathogen Moonlighting Proteins: From Ancestral Key Metabolic Enzymes to Virulence Factors. Microorganisms. 2021 Jun 15;9(6): 1300. doi: 10.3390/microorganisms9061300.

40. Figueiredo BC, Da’dara AA, Oliveira SC, Skelly PJ. Schistosomes Enhance Plasminogen Activation: The Role of Tegumental Enolase. PLoS Pathog. 2015 Dec 11;11(12): e1005335. doi: 10.1371/journal.ppat.1005335.

41. Jiang P, Zao YJ, Yan SW, Song YY, Yang DM, Dai LY, et al. Molecular characterization of a *Trichinella spiralis* enolase and its interaction with the host’s plasminogen. Vet Res. 2019 Dec 5;50(1): 106. doi: 10.1186/s13567-019-0727-y.

42. González-Miguel J, Morchón R, Siles-Lucas M, Oleaga A, Simón F. Surface-displayed glyceraldehyde 3-phosphate dehydrogenase and galectin from *Dirofilaria immitis* enhance the activation of the fibrinolytic system of the host. Acta Trop. 2015 May;145: 8–16. doi: 10.1016/j.actatropica.2015.01.010.

43. Boukli NM, Delgado B, Ricaurte M, Espino AM. *Fasciola hepatica* and *Schistosoma mansoni*: identification of common proteins by comparative proteomic analysis. J Parasitol. 2011 Oct;97(5): 852–861. doi: 10.1645/GE-2495.1.

44. Ravidà A, Cwiklinski K, Aldridge AM, Clarke P, Thompson R, Gerlach JQ, et al. *Fasciola hepatica* Surface Tegument: Glycoproteins at the Interface of Parasite and Host. Mol Cell Proteomics. 2016 Oct;15(10): 3139–3153. doi: 10.1074/mcp.M116.059774.

45. Morphew RM, Wright HA, LaCourse EJ, Woods DJ, Brophy PM. Comparative proteomics of excretory-secretory proteins released by the liver fluke *Fasciola hepatica* in sheep host bile and during in vitro culture ex host. Mol Cell Proteomics. 2007 Jun;6(6): 963–972. doi: 10.1074/mcp.M600375-MCP200.

46. Hanna RE. *Fasciola hepatica*: glycocalyx replacement in the juvenile as a possible mechanism for protection against host immunity. Exp Parasitol. 1980 Aug;50(1): 103–114. doi: 10.1016/0014-4894(80)90012-0.

47. Marcilla A, Trelis M, Cortés A, Sotillo J, Cantalapiedra F, Minguez MT, et al. Extracellular vesicles from parasitic helminths contain specific excretory/secretory proteins and are internalized in intestinal host cells. PLoS One. 2012;7(9): e45974. doi: 10.1371/journal.pone.0045974.

48. Cwiklinski K, de la Torre-Escudero E, Trelis M, Bernal D, Dufresne PJ, Brennan GP, et al. The Extracellular Vesicles of the Helminth Pathogen, *Fasciola hepatica*: Biogenesis Pathways and Cargo Molecules Involved in Parasite Pathogenesis. Mol Cell Proteomics. 2015 Dec;14(12): 3258–3273. doi: 10.1074/mcp.M115.053934.

49. Trelis M, Sánchez-López CM, Sánchez-Palencia LF, Ramírez-Toledo V, Marcilla A, Bernal D. Proteomic Analysis of Extracellular Vesicles From *Fasciola hepatica* Hatching Eggs and Juveniles in Culture. Front Cell Infect Microbiol. 2022 Jun 3;12: 903602. doi: 10.3389/fcimb.2022.903602.

50. Sánchez-López CM, González-Arce A, Soler C, Ramírez-Toledo V, Trelis M, Bernal D, et al. Extracellular vesicles from the trematodes *Fasciola hepatica* and *Dicrocoelium dendriticum* trigger different responses in human THP-1 macrophages. J Extracell Vesicles. 2023 Apr;12(4): e12317. doi: 10.1002/jev2.12317.

51. Ubeira FM, Martínez-Sernández V, González-Warleta M, Mezo M. Diagnostics for Animal and Human Fasciolosis. In: Dalton JP, editor. Fasciolosis. Wallingford, UK: CAB International; 2022. pp. 438-479.

52. Morales A, Espino AM. Evaluation and characterization of *Fasciola hepatica* tegument protein extract for serodiagnosis of human fascioliasis. Clin Vaccine Immunol. 2012 Nov;19(11):1870–1878. doi: 10.1128/CVI.00487-12.

53. Celik F, Simsek S, Selcuk MA, Kesik HK, Gunyakti Kilinc S, Aslan Celik B. Cloning and expression of *Fasciola hepatica* enolase gene and efficacy of recombinant protein in the serodiagnosis of sheep fasciolosis. Vet Parasitol. 2023 Aug;320: 109961. doi: 10.1016/j.vetpar.2023.109961.

54. Piast M, Kustrzeba-Wójcicka I, Matusiewicz M, Banaś T. Molecular evolution of enolase. Acta Biochim Pol. 2005;52(2): 507–513.

55. Mahana N, Abd-Allah HA, Salah M, Tallima H, El Ridi R. *Fasciola gigantica* enolase is a major component of worm tegumental fraction protective against sheep fasciolosis. Acta Trop. 2016 Jun;158: 189–196. doi: 10.1016/j.actatropica.2016.03.009.

56. Chen N, Yuan ZG, Xu MJ, Zhou DH, Zhang XX, Zhang YZ, et al. *Ascaris suum* enolase is a potential vaccine candidate against ascariasis. Vaccine. 2012 May 14;30(23): 3478–3482. doi: 10.1016/j.vaccine.2012.02.075.

57. Wang X, Chen W, Tian Y, Mao Q, Lv X, Shang M, et al. Surface display of *Clonorchis sinensis* enolase on *Bacillus subtilis* spores potentializes an oral vaccine candidate. Vaccine. 2014 Mar 10;32(12):1338–1345. doi: 10.1016/j.vaccine.2014.01.039.

58. Zhang S, Guo A, Zhu X, You Y, Hou J, Wang Q, et al. Identification and functional characterization of alpha-enolase from *Taenia pisiformis* metacestode. Acta Trop. 2015 Apr;144: 31–40. doi: 10.1016/j.actatropica.2015.01.007.

59. Zhang X, Xu L, Song X, Li X, Yan R. Molecular cloning of enolase from *Trichinella spiralis* and the protective immunity in mice. Acta Parasitol. 2018 Jun 26;63(2): 252–260. doi: 10.1515/ap-2018-0029.

60. Wistow GJ, Lietman T, Williams LA, Stapel SO, de Jong WW, Horwitz J, et al. Tau-crystallin/alpha-enolase: one gene encodes both an enzyme and a lens structural protein. J Cell Biol. 1988 Dec;107(6 Pt 2): 2729-2736. doi: 10.1083/jcb.107.6.2729.

61. Kainulainen V, Korhonen TK. Dancing to another tune-adhesive moonlighting proteins in bacteria. Biology (Basel). 2014 Mar 10;3(1): 178–204. doi: 10.3390/biology3010178.

62. Allu I, Sahi AK, Koppadi M, Gundu S, Sionkowska A. Decellularization Techniques for Tissue Engineering: Towards Replicating Native Extracellular Matrix Architecture in Liver Regeneration. J Funct Biomater. 2023 Oct 16;14(10): 518. doi: 10.3390/jfb14100518.

63. Plow EF, Herren T, Redlitz A, Miles LA, Hoover-Plow JL. The cell biology of the plasminogen system. FASEB J. 1995 Jul;9(10): 939–945. doi: 10.1096/fasebj.9.10.7615163.

64. Bernal D, de la Rubia JE, Carrasco-Abad AM, Toledo R, Mas-Coma S, Marcilla A. Identification of enolase as a plasminogen-binding protein in excretory-secretory products of *Fasciola hepatica*. FEBS Lett. 2004 Apr 9;563(1-3): 203–206. doi: 10.1016/S0014-5793(04)00306-0.

65. González-Miguel J, Valero MA, Reguera-Gomez M, Mas-Bargues C, Bargues MD, Simón F, et al. Numerous *Fasciola* plasminogen-binding proteins may underlie blood-brain barrier leakage and explain neurological disorder complexity and heterogeneity in the acute and chronic phases of human fascioliasis. Parasitology. 2019 Mar;146(3): 284–298. doi: 10.1017/S0031182018001464.

66. Lijnen HR. Elements of the fibrinolytic system. Ann N Y Acad Sci. 2001;936: 226–236. doi: 10.1111/j.1749-6632.2001.tb03511.x.

